# MassBlast: A workflow to accelerate RNA-seq and DNA database analysis

**DOI:** 10.1101/131953

**Authors:** André Veríssimo, Jean-Etienne Bassard, Alice Julien-Laferrière, Marie-France Sagot, Susana Vinga

**Affiliations:** IDMEC, Instituto Superior Técnico, Universidade de Lisboa, Av. Rovisco Pais, 1049-001 Lisboa, Portugal; INRIA Grenoble Rhône-Alpes & Laboratoire de Biométrie et Biologie Évolutive, Université Claude Bernard (Lyon 1); Copenhagen Plant Science Center, Plant Biochemistry Laboratory, Department of Plant and Environmental Science, University of Copenhagen

**Author notes:** Equal contributors (André Veríssimo, Jean-Etienne Bassard).

## Abstract

**Summary:** Current workflows for sequence analysis heavily depend on user input and manual curation. New specialized tools and methods are appearing all the time, but the actions required for a full analysis are disconnected and very time-consuming. The software we propose, MassBlast, combines BLAST+ and an automated workflow analysis to filter the results and significantly improve the annotation of multiple sequencing databases for exploring new biosynthetic pathways and new protein families, among other applications. MassBlast is fully configurable and reproducible.

**Availability and Implementation:** The MassBlast package is written in Ruby. Source code and releases are freely available from Github (https://github.com/averissimo/mass-blast) for all major platforms (Linux, MS Windows and OS X) under the GPLv3 license.

## 1 Introduction

The increasing availability of sequencing data is accompanied by many computational challenges for efficient analysis and querying of large databases of sequences. In fact, these tasks are still highly dependent on demanding pre-existing manual curation efforts, which involve managing large data sets while processing and combining results derived from using different analytical tools. A key task which requires optimization is the search for gene sets associated with specific pathways or gene families. This task is generally undertaken by searching for homologous coding sequences in sequencing databases, followed by curation of the results to identify, select and annotate the best matches. Tools such as Blast2Go (Conesa *et al.*, 2005) and Multblast (Mittler *et al.*, 2010) can annotate the transcriptome, but they lack curation functionalities that are as important as the primary homology-based identification of candidate genes. The goal of our research is to improve and accelerate annotation of sequencing databases by developing a pipeline with user-defined parameters that allows operators to query multiple databases for matches to target sequences and to identify rapidly the best candidates. Our new tool, hereafter referred to as MassBlast, allows the best alignment of pairs of sequences between a dataset consisting of contigs obtained from sequencing assemblies and fasta files with known sequences, using parameters specified by the user.

## 2 Implementation

MassBlast combines multiple tools in a single automated workflow (Fig. 1), where process management and tool execution are implemented in the Ruby programming language. Data are analysed sequentially by: (1) performing BLAST (Altschul *et al.*, 1990) queries on sequencing data with the goal of identifying whether target genes are being represented; (2) combining all BLAST results into a single file that can be analysed separately; (3) discarding all results outside a configurable identity range, with upper and lower limits defined by the user; (4) filtering out all the redundant BLAST results as defined by user-specified parameters; (5) finding the longest Open Read Frame (ORF) in the candidate sequences using the ORF Finder library (https://github.com/averissimo/orf_finder), and saving the resulting ORF in a fasta file usable by other bioinformatic tools.

**Fig. 1.**
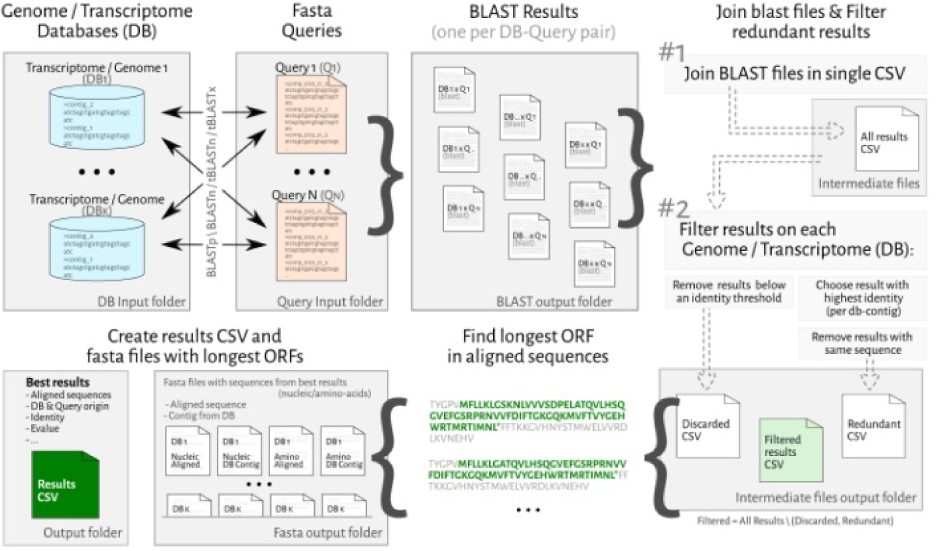
MassBlast workflow.

There are three BLAST search methods that are currently supported in MassBlast, each adapted for different combinations of both queries and databases:

- Blastn: Nucleic-acid sequences against a nucleic-acid database.
- tBlastn: Protein sequences against a nucleic-acid database (dynamically translated to amino-acid sequences in all six reading frames).
- tBlastx: Nucleic-acid sequences against nucleic-acid database, where both query and database are dynamically translated to amino-acid sequences into all six reading frames.

The user can tailor the workflow to specific uses by changing the configuration of each step. For example, the filtering step can be customized. Although it is designed to find the homologous results within an interval of similarity (by default from 40% to 100%), filtering can also be used to return the worst results of an interval, which may be of interest when searching for partially similar sequences or new unnamed sequences. Other customizable options include the BLAST parameters, the similarity criteria for discarding hits, how to find redundant hits, the codon preference table to be used to translate aligned sequences, the start and stop codon definitions when searching for the longest ORFs, among all options described fully in the online operating manual.

All resulting nucleotides and amino-acid sequences are saved in fasta to allow further use with other bioinformatics software.

MassBlast requires BLAST+ and a Ruby environment with bioinformatic libraries, Bioruby (Goto *et al.*, 2010) and ORF Finder. The environment can either be manually installed by the user or be deployed by a release package (https://github.com/averissimo/mass-blast/releases) that already contains the Ruby interpreter and all Ruby libraries. All options come pre-configured with the packaged version of MassBlast, allowing the user to rapidly run the application with its default workflow settings. Once installed, the next step requires the user to copy the fasta query files and databases into the respective folders and to run the workflow by using the distributed executable: mass-blast.

The workflow can be tested on the Blackberry transcriptome (Garcia-Seco *et al.*, 2015) together with benchmarked queries (cytochromes P450 from the CYP71 clan), which test the different search methods and postprocessing filtering steps.

## 3 Application

The analysis of nucleotide datasets is very complex and relies on an array of tools and processes for retrieving relevant information. Automated workflows such as MassBlast can play an important role to accelerate data analysis. Although MassBlast does not offer a novel concept for data analysis, it represents a new pipeline that allows the rapid assessment of data in the context of different hypotheses.

As a test example, when analyzing the Blackberry transcriptome comprising 42431 contigs against 50 cytochromes P450 from the CYP71 clan, it took 6h42min for a trained scientist to do the job manually. Manual data mining was carried out using the BLAST engine implemented in BioEdit, followed by a manual curation and search for ORFs with online tools. In contrast, it took 6s using MassBlast with a personal computer (1 core of an intel(R) i7 4980HQ @2.8Ghz processor, 32 GB DDR3L RAM, SSD drive, windows 10 operating system) to replicate the manual results. During the MassBlast execution, 63% of the computing time was used for the BLAST step. The latter is heavily dependent on the size of the databases and queries, whereas the curation phase depends exclusively on the results from BLAST. Most of the processing time is spent loading the data from disk. Using the MassBlast workflow, the amount of effort required for analysis and validation has been greatly reduced, because MassBlast not only accelerates these processes, but also provides reproducible results every time a query is run.

## 4 Conclusion

MassBlast can BLAST several query files against several databases. MassBlast can be used to search and/or annotate sequences (*e.g.* specific biosynthetic pathways, all cytochromes P450, etc.), to search new protein families (*e.g.* not yet named cytochromes P450), for a quick exploration of new RNAseq libraries, for a quick comparison of RNAseq libraries, or to generate lists of candidates that can then be given as input to other tools.

## Acknowledgements

We would like to thank Cathie Martin and Philippe Vain for reading the manuscript and providing us with important comments and insights. We would also like to thank: Aldo Ricardo Almeida Robles and Nuno Mira for testing MassBlast.

